# Analysis of the Genetic Structure and Diversity of Upland Cotton Groups in Different Planting Areas Based on SNP Markers

**DOI:** 10.1101/2021.06.11.448075

**Authors:** Jungduo Wang, Zeliang Zhang, Zhaolong Gong, Yajun Liang, Xiantao Ai, Zhiwei Sang, Jiangping Guo, Xueyuan Li, Juyun Zheng

**Author notes:** These authors have contributed equally to this work (Jungduo Wang, Zeliang Zhang, Zhaolong Gong).

## Abstract

Genetic diversity, kinship and population genetic structure analyses of *Gossypium hirsutum* germplasm can provide a better understanding of the origin and evolution of *G. hirsutum* biodiversity. In this study, 1313331 SNP molecular markers were used to construct a phylogenetic tree of each sample using MEGAX, to perform population structure analysis by ADMIXTURE software and principal component analysis (PCA) by EIGENSOFT software, and to estimate relatedness using SPAGeDi. ADMIXTURE software divided the experimental cotton population into 16 subgroups, and the *Gossypium hirsutum* samples could be roughly clustered according to source place, but there were some overlapping characteristics among samples. The experimental cotton population was divided into six groups according to source to calculate the genetic diversity index (*H*), and the obtained value (0.306) was close to that for germplasm collected by others in China. Cluster 4 had a relatively high genetic diversity level (0.390). The degrees of genetic differentiation within the experimental cotton population groups were low (the population differentiation indexes ranged from 0.02368 to 0.10664). The genetic distance among cotton accessions varied from 0.000332651 to 0.562664014, with an average of 0.25240429. The results of this study may provide a basis for mining elite alleles and using them for subsequent association analysis.

## Introduction

Cotton is one of the most important economic crops and a significant component of Chinese economy; moreover, it is widely cultivated worldwide and has a long history. It is the main textile material in the global textile industry, accounting for more than 60% of raw domestic textile materials and more than 50% of the total sales of fiber-based products in international consumer markets. Cotton is the foremost source of natural fiber, and cottonseed oil is considered a very-good-quality dietary fiber oil that accounts for approximately 10% of the global production of animal and plant oils. Cotton kernels are rich in protein and an important source of protein for humans; kernel powder from low-phenol cotton is also used as a major additive in high-grade foods. Cotton stalks can act as combustion materials and, after crushing, can provide crude feed for the livestock industry and can be used as feedstock in some industrial aspects (Liu.2015). With the increase in textile industrial development and improvements in the scientific and mechanistic levels of crop cultivation in China, the country has gradually become the largest cotton producer in the world, as well as the largest cotton consumer, with an annual planting of up to 80 million mu, providing the motivation for economic development and to meet the needs of the people.

In accordance with its botanical classification, cotton is a dicotyledon that belongs to the clade Angiospermae, order Malvales, family Malvaceae, and genus Gossypium. Beasley divided diploid cotton species into five groups, a to e, based on kinship and local environmental conditions, to lay a foundation for subsequent studies on the classification of cotton species (Beasley *et al*.1940). In 1978, Fryxell (1979) divided the genus Gossypium into four cultivated species and 39 other wild species based on previous works, and this classification remained partially controversial, even though it was mostly recognized. Afterwards, Fryxell (1992) summarized the genus Gossypium into 50 species. A few years ago, botanists again discovered two new tetraploid cotton species, *G. ekmanianumsi* and *G. stephensi* (Grover *et al*.2015;Gallagher *et al*.2017). Currently, the genus Gossypium can be divided into a total of 52 species, including seven allotetraploid species and 45 diploid species.

At present, some researchers use sequence-related amplified polymorphisms (SRAPs), simple sequence repeats (SSRs), amplified fragment length polymorphisms (AFLPs), single-nucleotide polymorphisms (SNPs) and other molecular markers to study cotton genetic diversity. Dong (2007) used SSR markers to evaluate the diversity of 96 germplasm resources of upland cotton, sea island cotton, Asian cotton and grass cotton and found that the main phenotypic traits of cotton were significantly different between germplasms and within species. An extremely significant finding was that the phenotypic diversity index of upland cotton germplasm was the highest. Wu *et al*. (2001) studied 36 varieties using ISSR(inter-simple sequence repeat) technology and showed that the hereditary basis of upland cotton cultivars is relatively narrow. Liu *et al*. (2003) used RAPD marker technology to analyze the genetic diversity of 166 representative cotton varieties (or lines) in China since the founding of the People’s Republic and showed that the genetic range of selfing upland cotton varieties in China was narrower than that of imported varieties. The hereditary basis of hybrid upland cotton is narrower than that of the conventional variety; that of the upland cotton varieties generated after the 1980s is narrower than that of the varieties from the 1970s; that of the Yangtze River cotton varieties is narrower than that of the Huanghuai cotton varieties; and that of the northwest inland cotton varieties is narrower than that of the Yangtze River cotton varieties. Multani *et al*. (1995) used RAPD technology to analyze 14 Australian cotton varieties and found that their genetic relationships were relatively close. Cotton also shows a certain degree of differentiation at the molecular level. Gao *et al*. (2010) used SSR technology to analyze the genetic diversity of tetraploid cotton species in China. The results showed that wild lines of broad-leaved cotton and upland cotton have the closest genetic relationship, and the genetic relationship between Darwin cotton and sea island cotton was very close. Brown cotton was also relatively closely related to these types of cotton. Wu (2012) analyzed the diversity of 168 sea island cotton samples by morphological observation combined with SRAP technology, which showed that the genetic basis of Chinese sea island cotton is narrow and that the level of diversity is low. Kuang *et al*. (2011) used 36 pairs of primers with high polymorphism to analyze the genetic diversity of 32 main cultivated varieties selected in 2008 in China. Through cluster analysis, they found that the genetic differences among cotton varieties in the Yangtze River Basin were the largest, the differences in the Xinjiang cotton area were the second largest, and the differences in the Yellow River basin were the smallest. The genetic diversity of hybrids is richer than that of conventional species. Chen *et al*. (2006) analyzed the diversity of 43 upland cotton basic germplasms in China and found that their genetic diversity level showed a downward trend, and the diversity level of cotton areas in the Yangtze River and Yellow River basins was lower than that of foreign areas where basic germplasms are grown. Thus, the genetic backgrounds of the breeds are relatively narrow.

Population genetics is a discipline in which mathematical and statistical methods are applied to study gene and genotype frequencies in populations and the effects of selection and mutations that influence these frequencies in order to study the relationships of processes such as migration and genetic drift with genetic structure, thereby exploring evolution. Analysis of genetic differences using molecular markers is an essential approach in population genetics. Population genetics studies are often performed by using SNP markers.

Before performing GWAS analysis, clarifying the genetic background and population structure of the tested material is fundamental. Therefore, the work carried out in this paper will lay a foundation for further association analysis.

## 1 Materials and methods

### 1.1 Experimental materials

This study used 273 domestic and foreign resources of upland cotton varieties (Appendix 1) as research materials, and all materials were provided by the Economic Crop Research Institute of the Xinjiang Academy of Agricultural Sciences. Middle intact leaves were collected from individual plants grown in the field.

### 1.2 Extraction and sequencing of cotton DNA

The genomic DNA of cotton leaves was extracted by the modified CTAB method and then tested. Enzymatic 3’ A processing, attachment of a dual-index (Kozich *et al*.2013) sequencing adapter, PCR amplification, purification, mixing, and sequencing on an Illumina sequencing platform were performed. Japanese rice was used as a sequencing control to evaluate the accuracy of the enzymatic experiment. The dual-index adapter was used to identify the original data obtained by sequencing and obtain the reads from each sample. After filtering the adapters from the sequencing reads, sequencing quality and data volume assessments were performed. The digestion efficiency of HaeIII+SspI-HF® was evaluated through Nipponbare rice data to judge the accuracy and effectiveness of the experimental process.

### 1.3 Development of cotton snp markers

The sequenced reads obtained by simplified sequencing needed to be realigned to the reference genome to perform subsequent variation analysis. Using Zhejiang University Cotton v2.1 (download:https://www.cottongen.org/data/download/genome_tetraploid/TM-1) as the reference genome, BWA 0.7.15 (Li *et al*.2009) was used to compare sequenced reads to the reference genome. Using GATK 4.0 (McKenna *et al*.2010) and SAMtools 1.9 (Li *et al*.2009b) methods, SNPs and SNP marker intersections were identified to create the final reliable SNP marker dataset. SnpEff 4.0 (Cingolani *et al*.2012) software was used to obtain the locations of the variable sites (intergenic zones, gene zones, or CDS zones) in the reference genome and the effects of the variations (synonymous mutations, nonsynonymous mutations, etc.).

### 1.4 Analysis of cotton genetic evolution

#### 1.4.1 Phylogenetic analysis

A phylogenetic tree is used to indicate the evolutionary relationships between species that are considered to have a common ancestor and to describe the classification and evolutionary relationships between species. In the analysis, according to the genetic data of the population, the distance of the genetic relationship between the materials was inferred, a distance matrix was constructed, and a phylogenetic tree was created based on the distance matrix. MEGAX 7.0.14 software was used to construct the phylogenetic tree of each sample based on the neighbor-joining method and the p-distance model, and bootstrapping was repeated 1,000 times.

#### 1.4.2 PCA

Principal component analysis is a statistical method that is performed by transforming a set of correlated variables into a set of linearly uncorrelated variables; the transformed set of variables is called the principal components. In population genetics, different materials are clustered into different subgroups based on the degree of SNP differentiation (or degree of divergence) between materials, and the results can be used for mutual verification with the results of other clustering methods. Based on the SNPs identified, EIGENSOFT 7.2.1 (Alkes *et al*.2006) software was used to perform principal component analysis to cluster the samples. Through PCA, we could determine which samples were relatively closely related and which samples were relatively distantly related, facilitating evolutionary analysis.

#### 1.4.3 Analysis of genetic structure

Population structure, also known as group stratification, refers to the existence of subgroups with different gene frequencies in the studied group. The materials in the same subgroup are closely related, and the subgroups are relatively distantly related. Group structure analysis can reveal the number of ancestors of the studied group and the blood origin of each sample. The group cluster analysis method is currently widely used, as it helps to understand the evolution of materials. Based on the SNPs identified, ADMIXTURE V250 (Alexander *et al*. 2009) software was used to analyze the group structure of the research materials. For the research group, the number of subgroups (K value) was preset to 1-20 for clustering, the clustering results were cross-validated, and the optimal number of clusters was determined according to the lowest cross-validation error rate.

#### 1.4.4 Genetic relationship analysis

Estimation of the affinity (relative kinship) between natural groups can be performed using SPAGeDi 1.3 (Hardy *et al*.2002) software. The kinship itself is the relative value that defines the genetic similarity between two specific materials and that between any material and itself, so when the kinship value between the two materials is less than 0, it is directly defined as 0.

#### 1.4.5 Analysis of genetic diversity

Population genetic parameters and the population index (Fst) were calculated using VCFtools software (https://vcftools.github.io/index.html). According to Wright, when the group differentiation index (Fst) equals 0 or 1, there is no differentiation between subgroups or complete differentiation between subgroups, respectively. However, values of 0 < Fst < 0.05, 0.05, 0.05 ≤ FST <0.15, 0.15 ≤ FST <0.25, or 0.25 ≤ FST <1 indicate weak, moderate, strong or very strong genetic differentiation between subgroups, respectively.

## 2 Results and analysis

### 2.1 Genotype analysis of the cotton population

In the experiment, the digestion efficiency of HaeIII+SspI-HF® was 97.02%, and a total of 2,161.76 Mb reads were obtained. A total of 1,313,331 SNPs were obtained from the population through comparison with the cotton genome. The average Q30 of the sequences was 96.87%, and the average GC content was 36.50%. The Nipponbare rice sequencing used to evaluate the accuracy of the experimental database yielded 2.12 Mb reads of data(Appendix 2).

Plotting the distribution of SNPs on chromosomes revealed that SNPs were present in high density at both ends of the chromosomes and at low density in the juxtacentromeric region. The densities of SNPs on chromosomes A07, A13, D01, D05, and D09 were relatively high (Figure 1). Chromosome ends are enriched in functional genes, and the mid-centromere and near-centromere regions are mostly repetitive sequences. The density distribution of SNPs on chromosomes was in accordance with that expected.

**Figure 1.**
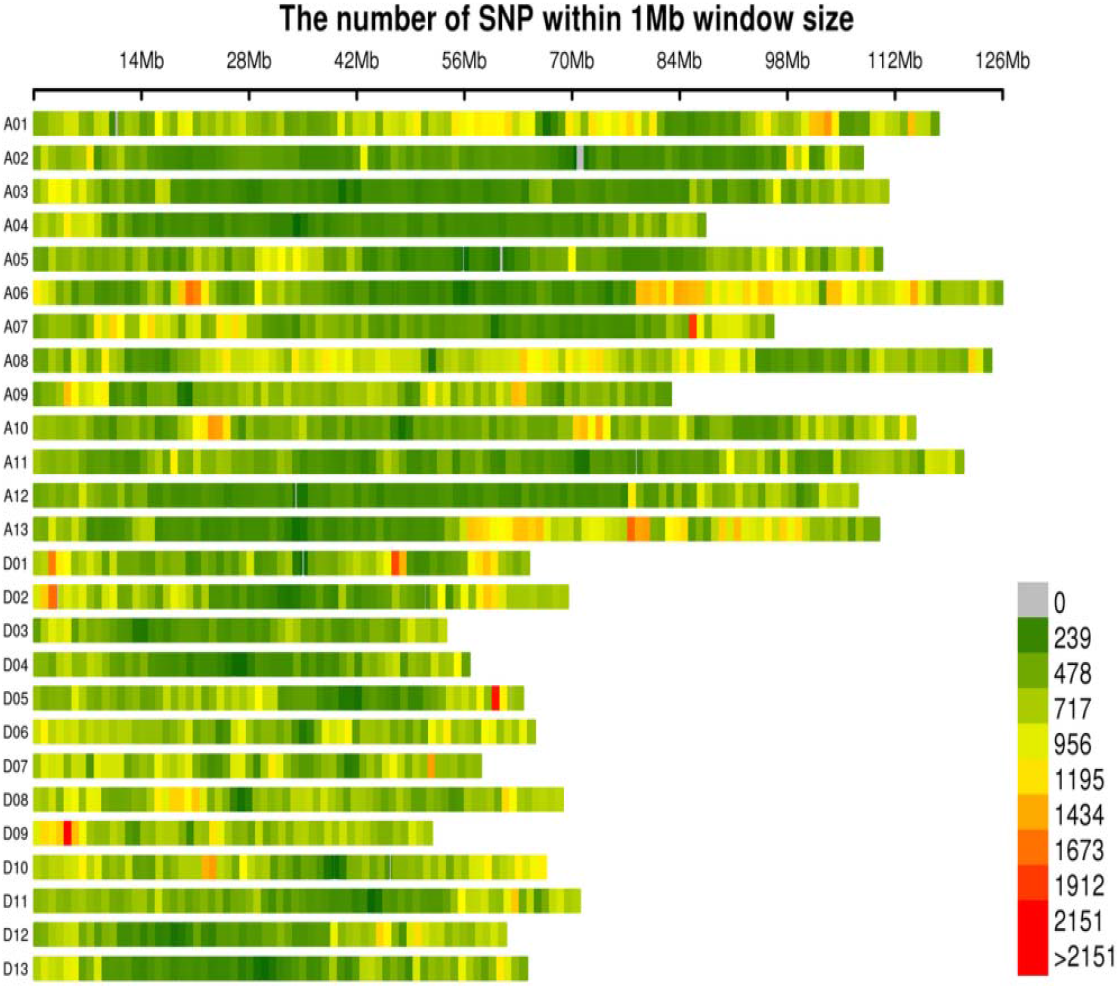
Distribution of SNPs on chromosomes.The abscissa is the length of the chromosome. Each band represents a chromosome. The genome is divided according to the size of 1Mb. The more SNP markers in each window, the darker the color, and the fewer SNP markers, and the lighter the color. ; The darker the color in the figure is the area where the SNP markers are concentrated.

### 2.2 Genetic evolutionary analysis

Phylogenetic trees were drawn based on the SNP markers, where most of the materials on the branches were from the inland cotton area of northwestern China, but they also contained materials from Central Asia(Figure 2).

**Figure 2.**
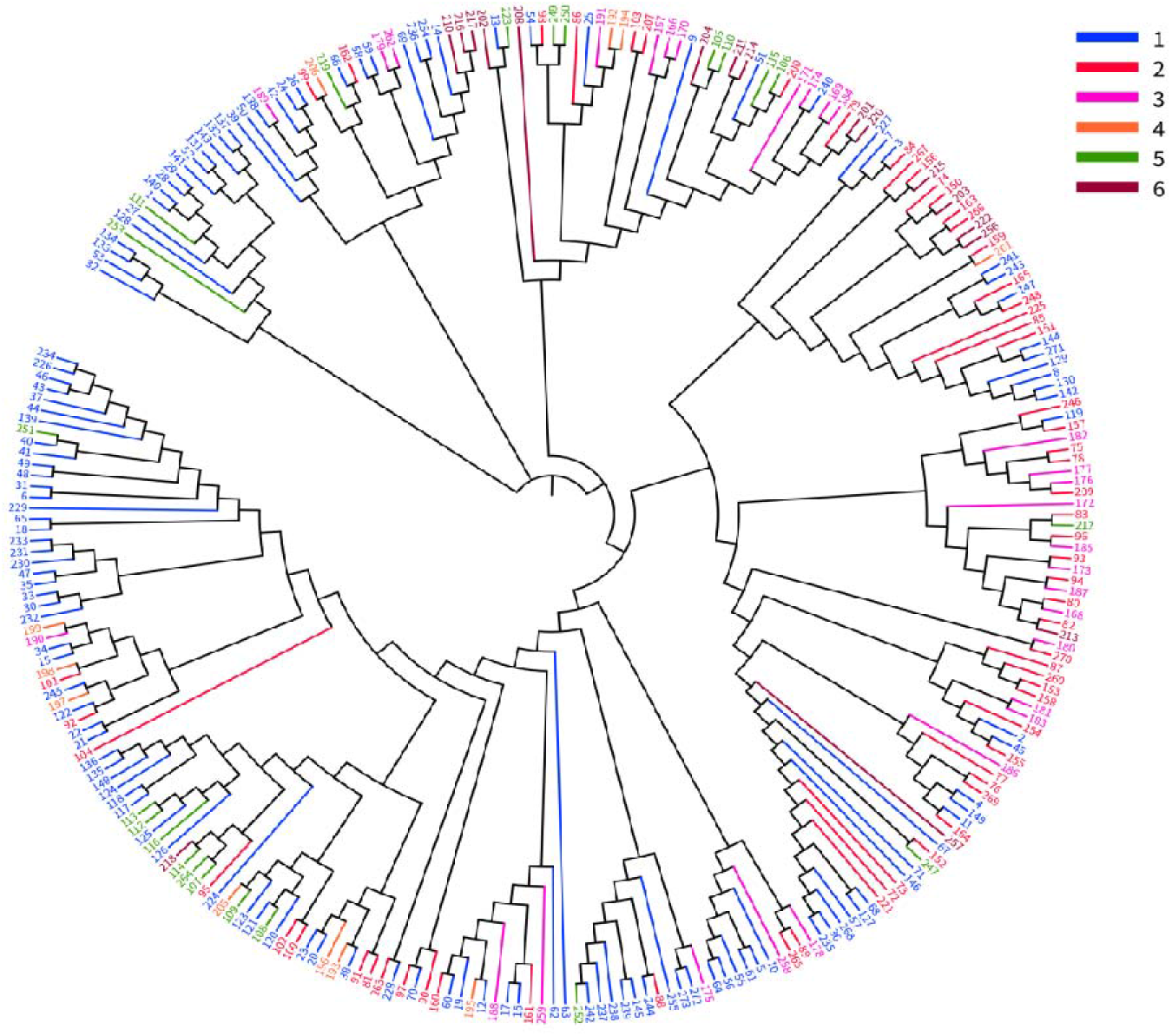
Phylogenetic tree.Each branch in the picture is a sample, and each source has a color. 1: Northwest Inland, 2: Yellow River Basin, 3: Yangtze River Basin, 4: Extra-early Cotton Area, 5: Central Asia, 6: United States and other sources.

Based on principal component analysis of 1,313,331 SNP molecular markers from the population, the first three components explained 11.15%, 8.26%, and 6.47% of the variation, respectively. The results of the first three components were used to draw a PCA plot in the R environment, which is shown in Figure 3. The cotton population showed a certain distribution gap, but most varieties were clustered together, with no obvious differences among the 273 cotton materials. In terms of group stratification, the varieties from the United States and Central Asia were close to the early Chinese land cotton varieties, which corresponds to the early introduction of Chinese land cotton varieties mainly from the United States and the Soviet Union and breeding with the introduced materials as the parents.

**Figure 3.**
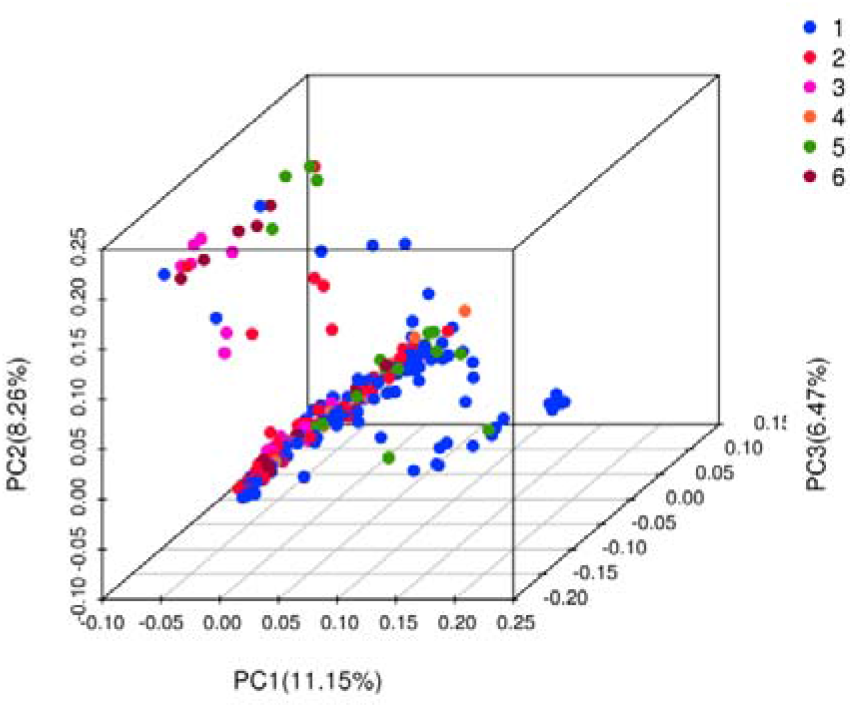
PCA three-dimensional clustering map. In the figure, the samples are gathered into three dimensions by PCA analysis, PC1 represents the first principal component, PC2 represents the second principal component; PC3 represents the third principal component. A point represents a sample, and a color represents a grouping 1: Northwest Inland, 2: Yellow River Basin, 3: Yangtze River Basin, 4: Extra-early Cotton Area, 5: Central Asia, 6: United States and other sources.

For the group genetic structure analysis of the 273 cotton varieties through 1,313,331 SNP molecular markers, the K value range for the number of subgroups was set to 1-20, and the cross-validation error (CV error) was calculated under the different K values. As shown in Figure 4, when K increased from 1 to 6, the CV error value decreased rapidly; when K increased from 6 to 11, the CV error value gradually decreased and tended to flatten; when K increased from 11 to 16, the CV error value showed a volatile, downward trend; and when K was greater than 16, the CV error value increased to a certain extent. Thus, when K was equal to 16, the CV error value is the smallest, and the 273 cotton variety groups could therefore be divided into 16 subgroups :Subgroup 1~Subgroup 16(Figure 4).

**Figure 4.**
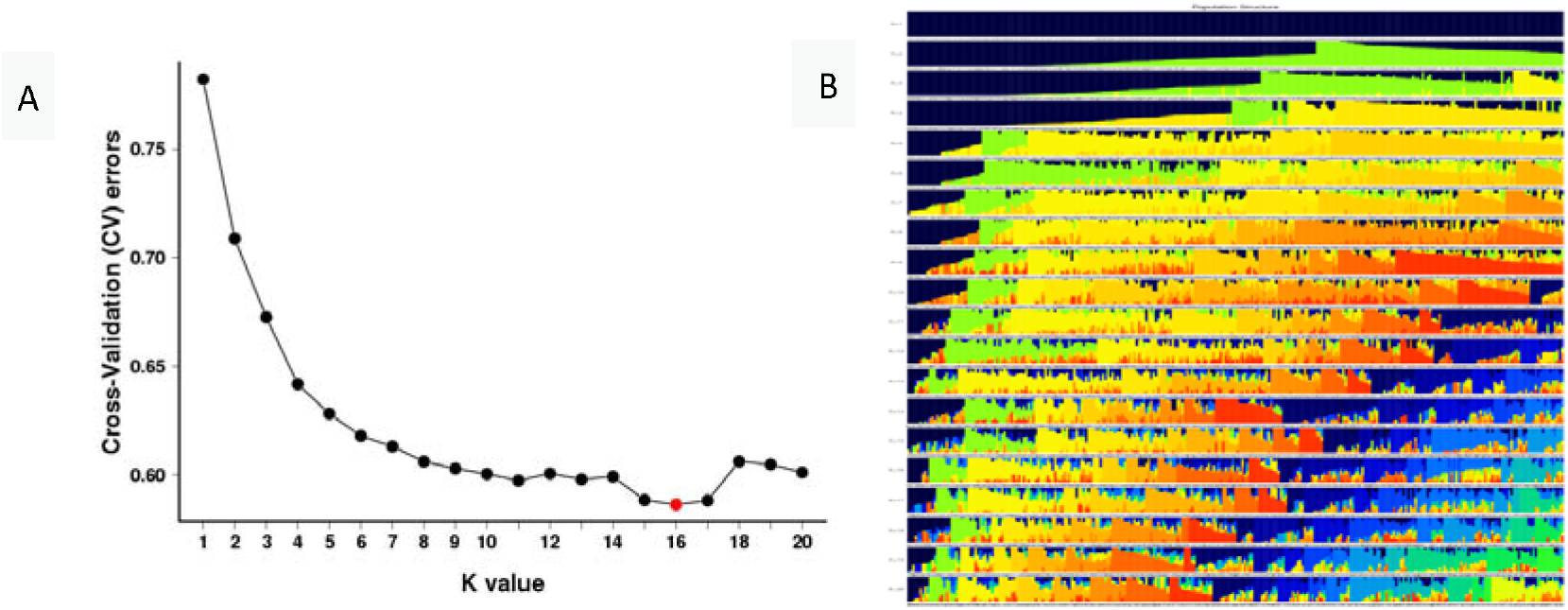
Cross-validation error rate of each K value of Admixture.(A) Cross-validation error rate for each K value, Group structure analysis of 273 cotton materials, calculated CV error. at K of 1~20. (B) Sample clustering results for each K value, Group structure at K =1-16 in which each individual is represented by lines of different colours, inferred which subgroup the breed belongs to from the proportion of color.

Depending on the Q value, each individual in the 16 subgroups was classified into the maximum-Q value subgroup (Table 1). The 16 subgroups were 9, 16, 52, 4, 10, 13, 8, 30, 13, 9, 26, 18, 8, 31, 5, and 21. Group structure analysis is more consistent with PCA and systematic evolutionary tree analysis, with independent tests on the subgroups and the source of the materials. The results of these analyses showed that the division of the subgroups was significantly correlated with the material source, indicating that the genetic background of the resources was relatively homogeneous.

**Table 1.** Specific species grouping

| Grouping | Variety number | Quantity |
| --- | --- | --- |
| Q1 | 1,29,37,43,44,46,139,226,234 | 9 |
| Q2 | 2,36,57,67,68,71,72,73,127,146,152,221,235,247,257,268 | 16 |
| Q3 | 10,13,25,58,59,63,66,75,78,80,82,83,87,89,94,96,99,119,153,154,155,157,158,162,168,172,173,176,177,178,179,180,181,182,183,185,186,187,206,209,212,213,219,223,246,258,259,260,262,265,267,270 | 52 |
| Q4 | 32,53,133,134 | 4 |
| Q5 | 4,8,11,76,77,130,142,149,164,269 | 10 |
| Q6 | 5,7,27,28,52,111,124,128,131,132,137,141,143 | 13 |
| Q7 | 55,56,61,64,147,165,241,243 | 8 |
| Q8 | 6,15,31,34,65,81,86,91,92,97,100,101,102,104,108,109,113,116,120,121,122,123,163,190,198,199,205,230,231,233 | 30 |
| Q9 | 14,24,26,39,42,69,138,189,210,216,217,236,254 | 13 |
| Q10 | 18,40,41,48,49,229,232,250,251 | 9 |
| Q11 | 20,23,35,38,47,50,70,90,95,107,112,114,117,118,125,135,136,140,148,193,196,202,224,228,263,264 | 26 |
| Q12 | 12,16,17,19,21,22,30,33,60,62,88,160,161,188,191,195,197,245 | 18 |
| Q13 | 145,237,238,239,242,244,252,255 | 8 |
| Q14 | 3,45,54,74,84,85,98,129,144,150,151,156,159,175,192,194,203,208,215,222,225,227,248,249,253,256,261,266,271,272,273 | 31 |
| Q15 | 106,115,126,200,218 | 5 |
| Q16 | 9,51,79,93,103,105,110,166,167,169,170,171,174,184,201,204,207,211,214,220,240 | 21 |

Kinship calculations were performed by TASSEL 3.0 software, and a heat map was generated in R (Figure 5), revealing that the 273 cotton varieties were closely related to most varieties (blue) and a few other materials (red).

**Figure 5.**
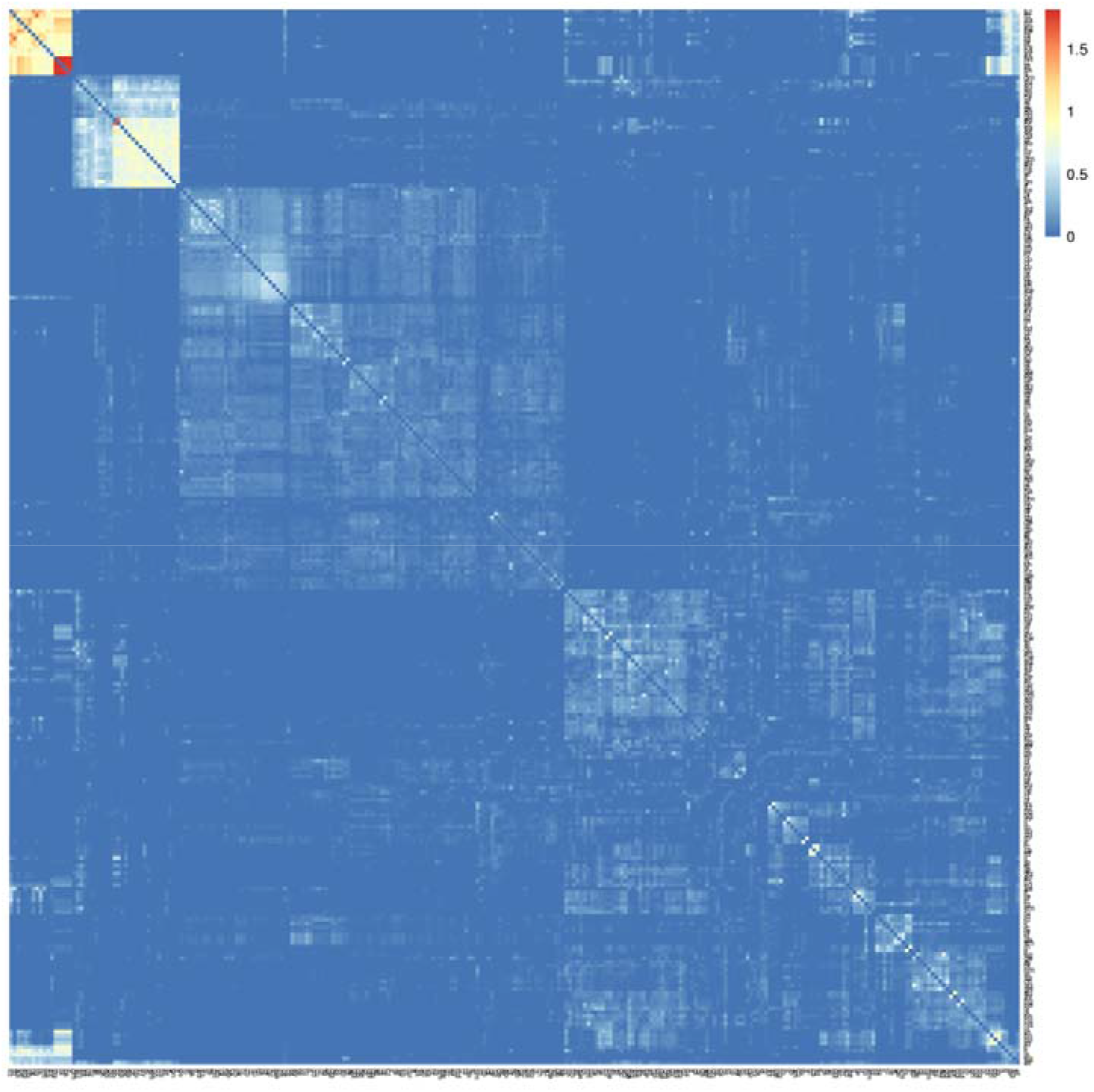
Heat map distribution map of genetic relationship

### 2.3 Analysis of cotton genetic diversity

The experimental cotton group was divided into 6 subgroups according to source, and 1,313,331 high-quality SNPs were evaluated to determine the genetic diversity of the experimental cotton group by calculating the diversity index, flavor index, and PIC. Through VCFtools calculations, the diversity index of the cotton population was determined to be 0.306, ranging between 0.314 for groups 1~6 and ~0.390, where group 2 had the lowest genetic diversity index (0.314) and group 4 had the highest (0.390) of the 6 groups. The aroma index (0.551) and PIC (0.296) were also determined. Group 4 had a relatively high genetic diversity level, but the overall genetic diversity level was still relatively low, which is consistent with the group structure analysis results(Table 2).

**Table 2.**
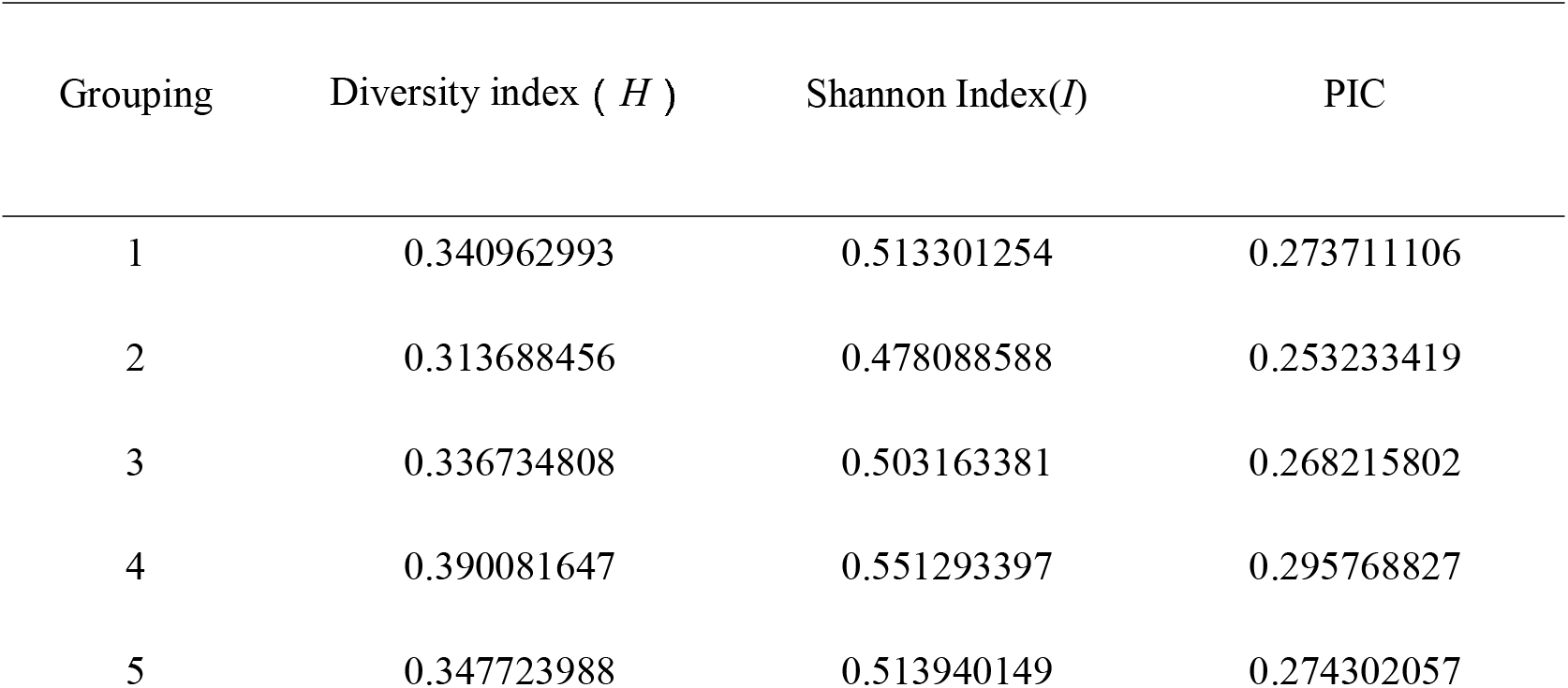
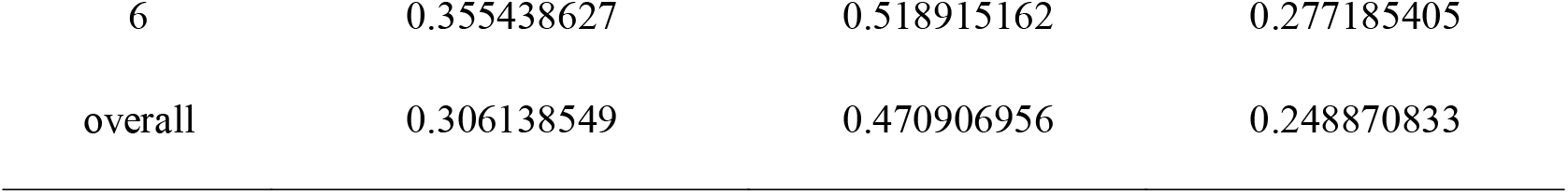
Statistics of genetic diversity

The population differentiation index (Fst) was used to evaluate the degree of differentiation between cotton groups and revealed that the groups were all moderately or weakly genetically differentiated, with genetic differentiation indexes between groups ranging between 0.02368 and 0.10664(Table 3). Among the groups, there was moderate differentiation between group 1 and groups 3, 4, and 5; there was moderate differentiation between group 2 and groups 3 and 4; there was moderate differentiation between groups 3, 4, 5 and 6; and the remaining groups were weakly equally divided. These results showed a moderate or weak degree of genetic differentiation among the groups; that is, the genetic relationship between the groups was relatively close. The genetic difference between group 3 and group 6 was the largest, and the difference between group 3 and group 5 was the smallest. This is related to the more effective promotion of selfing cotton varieties in the cotton planting area of the Yangtze River Basin. In addition, the genetic distance analysis of the cotton population used in this study showed that the genetic distance between these cotton germplasms ranged from 0.000332651~0.562664014. The average genetic distance was 0.25240429, and the genetic distances were quite different, indicating that there were large differences in gene exchange between subgroups. The genetic distance between Wanmian 8407 and Keyuan 1 was the closest (0.000332651), while that between Xinluzhong 4 and Ekangmian 33 was the longest (0.562664014).

**Table 3.**
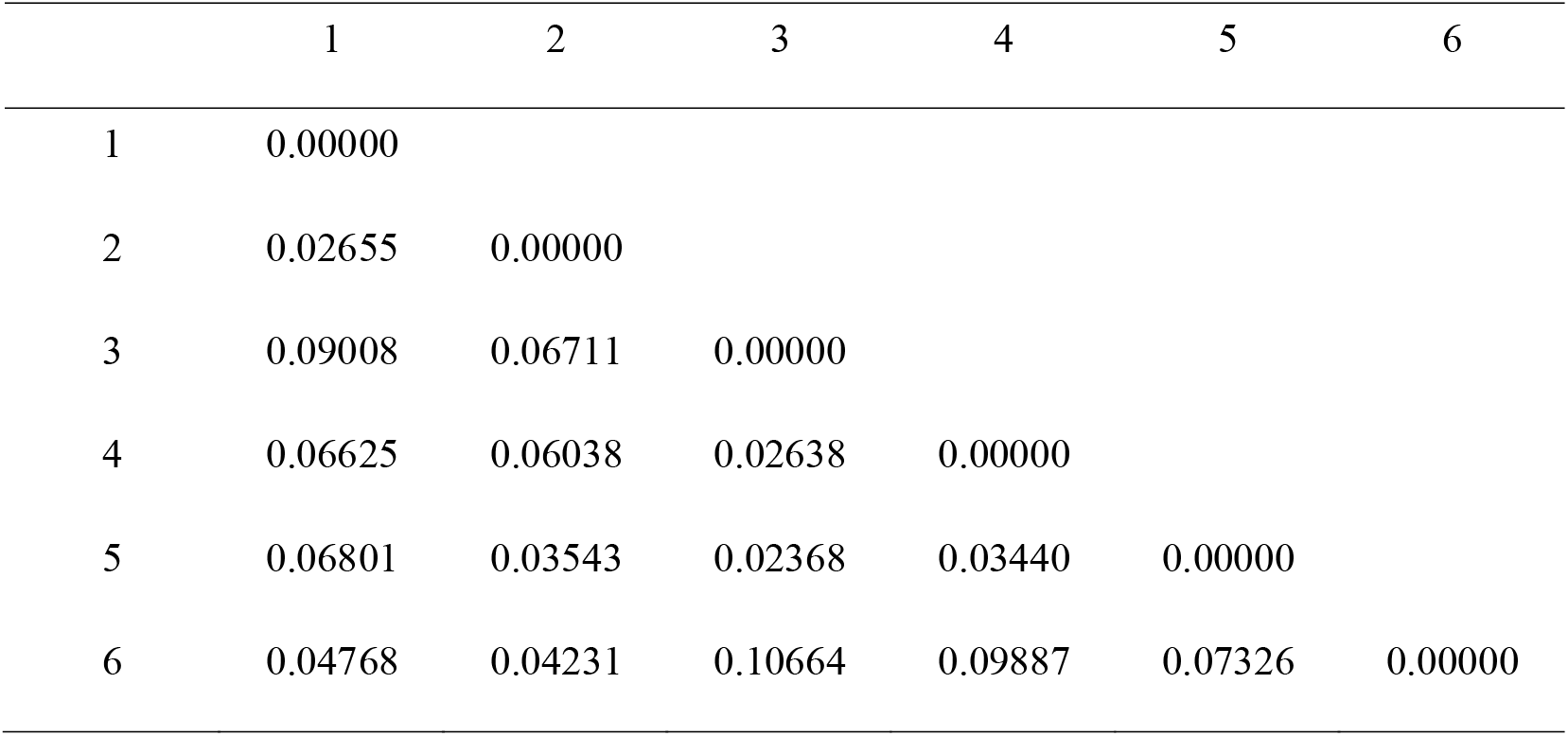
Statistics of population differentiation index

|  | 1 | 2 | 3 | 4 | 5 | 6 |
| --- | --- | --- | --- | --- | --- | --- |
| 1 | 0.00000 |  |  |  |  |  |
| 2 | 0.02655 | 0.00000 |  |  |  |  |
| 3 | 0.09008 | 0.06711 | 0.00000 |  |  |  |
| 4 | 0.06625 | 0.06038 | 0.02638 | 0.00000 |  |  |
| 5 | 0.06801 | 0.03543 | 0.02368 | 0.03440 | 0.00000 |  |
| 6 | 0.04768 | 0.04231 | 0.10664 | 0.09887 | 0.07326 | 0.00000 |

## 3 Discussion

### 3.1 Analysis of cotton population structure

Studies have shown that there will be false associations between genotypes and traits, which may be caused by population structure and an uneven distribution of alleles (Wu *et al*. 2019). To eliminate false positives in association analysis, we need to analyze the group structure of a test group first and control for that group structure. This study used ADMIXTURE software to analyze the population structure of 273 natural populations of cotton varieties at home and abroad. The results showed that when K=16, the CV error value was the smallest. Therefore, the 273 cotton germplasms were divided into 16 subgroups. According to the grouping results, in the same subgroup, most varieties were genetically related. The analysis results showed that some varieties with different origins were of one type, which may have been caused by crossover between varieties and environmental factors, but most varieties with the same origin and with similar genetic background information could be better classified. For subgroups, the results of population structure analysis were more in line with the evolutionary trends of the genetic background during breeding. According to the different origins of the experimental cotton materials, they were divided into 6 groups. The population differentiation indexes ranged between 0.02368 and 0.10664, and the population differentiation indexes between most groups were less than 0.07, indicating that the cotton population has moderately weak genetic differentiation. In the later period, the genetic relationship of varieties cultivated through continuous introduction and germplasm innovation gradually increased. China is not the origin of upland cotton. Previous studies have shown that since the introduction of tetraploid cotton to China, Daizimian 15 has been the most widely planted variety in China. In 1958 alone, more than 3.5 million hectares were planted, accounting for approximately 50% of the country’s cotton planting area that year. More than 100 varieties have been directly selected from Daizimian 15. In addition, there are quite a few varieties that are crossed with Daizimian 15. Principal component analysis and population structure analysis have also shown that the genetic range of cotton is relatively narrow, and most cotton varieties are derived from the same ancestor, Stormproof. The group differentiation of soybeans is driven by differences in geographical locations. Conversely, the formation of populations during the process of cotton breeding was mainly due to different ancestors, which is similar to the situation in wheat (Ye.2011;Mei.2012). There were some differences among the 33 local upland cotton varieties bred in southern Xinjiang collected by the Economic Crop Research Institute of the Xinjiang Academy of Agricultural Sciences, but the overall differences were not large; in cultivated cotton from other regions, the population differentiation indexes were between 0.01 and 0.05 (Ai *et al*.2010). This shows that cotton collected in China has a low degree of genetic differentiation. In comparison, the population differentiation index of cotton germplasm in this study is similar to that of cotton germplasm collected in China, and varieties from different ecological regions show obvious mixed characteristics.

### 3.2 Analysis of cotton genetic diversity

The sum of genetic information carried by all organisms is defined as genetic diversity, which can also be called species diversity. Genetic diversity is usually regarded as the sum of genetic variation among individuals within a species. In this study, the diversity index, Shannon index and PIC were calculated to evaluate the genetic diversity of the cotton population. Previous studies such as that of Ai (2017) genotyped 288 upland cotton germplasms; the genetic diversity was 0.31, and the PIC was 0.25. Fang *et al*. (2013) detected an average PIC of 0.29 in 193 upland cotton varieties collected from 26 countries. These results indicate that the diversity of cotton collected in China is relatively low and that the degree of genetic differentiation is low. Moreover, the level of genetic differentiation between landraces and modern cultivars is low (Tyagi *et al*. 2014). According to the cotton SNP molecular marker development and genetic diversity comparison, the genetic diversity index of each subgroup in the cotton population in this study was between 0.314 and 0.390, and the overall genetic diversity index was 0.306, with a PIC of 0.249. Thus, the genetic diversity index of the cotton germplasm is consistent with that of the cotton germplasm collected in China. The largest genetic diversity was observed between varieties from the the US and extra-early cotton regions, and the largest differentiation index was between varieties from the the US and the Yangtze River Basin. High-quality cotton varieties can be selected and introduced to China to enrich the existing cotton germplasm resources. In addition, the genetic relationship between the samples will also have a certain impact on the results of the association analysis. In this study, the genetic distances between 273 cotton germplasms were analyzed, and the kinship value was used to infer the genetic relationships between different materials. The results showed that the average genetic distance of these cotton germplasms was 0.252. Part of the results indicated that some varieties with different origins were clustered into one category. Varieties with similar genetic backgrounds can be better clustered into one category, and the clustering results are more in line with the evolutionary trends of the genetic background of the varieties.

## 4 Conclusion

Chinese cotton planting area is very large, but upland cotton was not domesticated in China. Upland cotton was first domesticated in the United States and was then introduced to China in the 1940s and 1950s. Among upland cotton varieties, Daizimian 15 and Sizimian 2B are the two most widely planted in China. On the basis of these two varieties, ,many modern cotton cultivars were bred by Chinese cotton breeders through genealogy and crossbreeding. After domestication and improvement, the yield of cotton was higher, the fiber quality was better, and the planting range was wider. Thus, these improved upland cotton cultivars can replace Asian cotton grown in China. The results of domestic and foreign studies have consistently shown that the genetic range of upland cotton varieties is narrow. The intraspecific genetic diversity was far lower than the interspecific differences (Liu *et al*. 2003;Curt *et al*. 1994). Using multiple methods to expand the genetic range of upland cotton varieties will be an important aspect of cotton germplasm resource innovation and breeding in the future. In this study, 1,313,331 SNP loci from the analyzed population were used to determine the population genetic structure of 273 domestic and foreign cotton germplasm resources. The results of the genetic diversity analysis revealed that the genetic diversity of the experimental cotton population was average; the results of the population genetic structure analysis showed that the population was divided into 16 subgroups (K=16). In addition, this cotton population was less differentiated than and closely related to the domestically collected cotton germplasm. Genetic distance analysis revealed that Wanmian 8407 is the closest genetically to Keyuan 1 and that Xinluzhong 4 is the most genetically distant from Ekangmian 33. The results of the genetic diversity analysis showed that the greatest genetic diversity occurred between cultivars from the US and early cotton regions, and the largest differentiation index was observed between cultivars from the US and the Yangtze River Valley. This study will provide a basis for genome-wide association analysis for mining elite genes and obtaining elite cotton germplasms.

## Dota availability statements

The raw sequence data reported in this paper have been deposited in the Genome Sequence Archive (Genomics, Proteomics & Bioinformatics 2017) in National Genomics Data Center (Nucleic Acids Res 2021), China National Center for Bioinformation / Beijing Institute of Genomics, Chinese Academy of Sciences, project number PRJCA 005438, under accession number CRAxxxxxx that are publicly accessible at https://ngdc.cncb.ac.cn/gsa.

## Acknowledge

We would like to thank Xinjiang Academy of Agricultural Sciences, China, for the cotton varieties provided for this study, BMK for the sequencing, and AJE for the English polishing. Thanks to all the people, units and enterprises who have provided help to this study.

## Funding

This work was supported by the National Natural Science Foundation of China (no. 31760405, U1903204).

## Conflicts of interest

The authors have no conflicts of interest to declare.

---

**Appendix 1.**
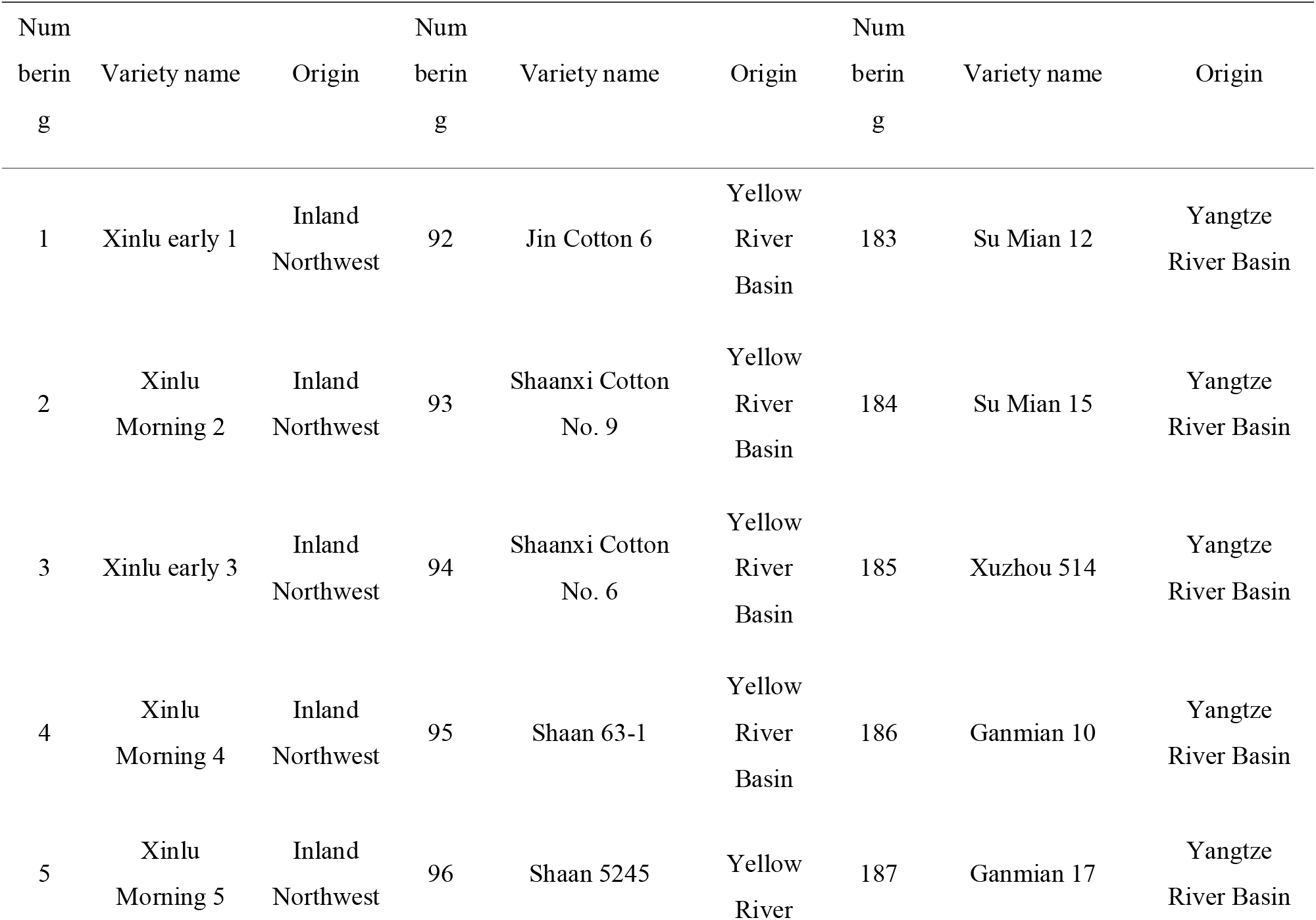
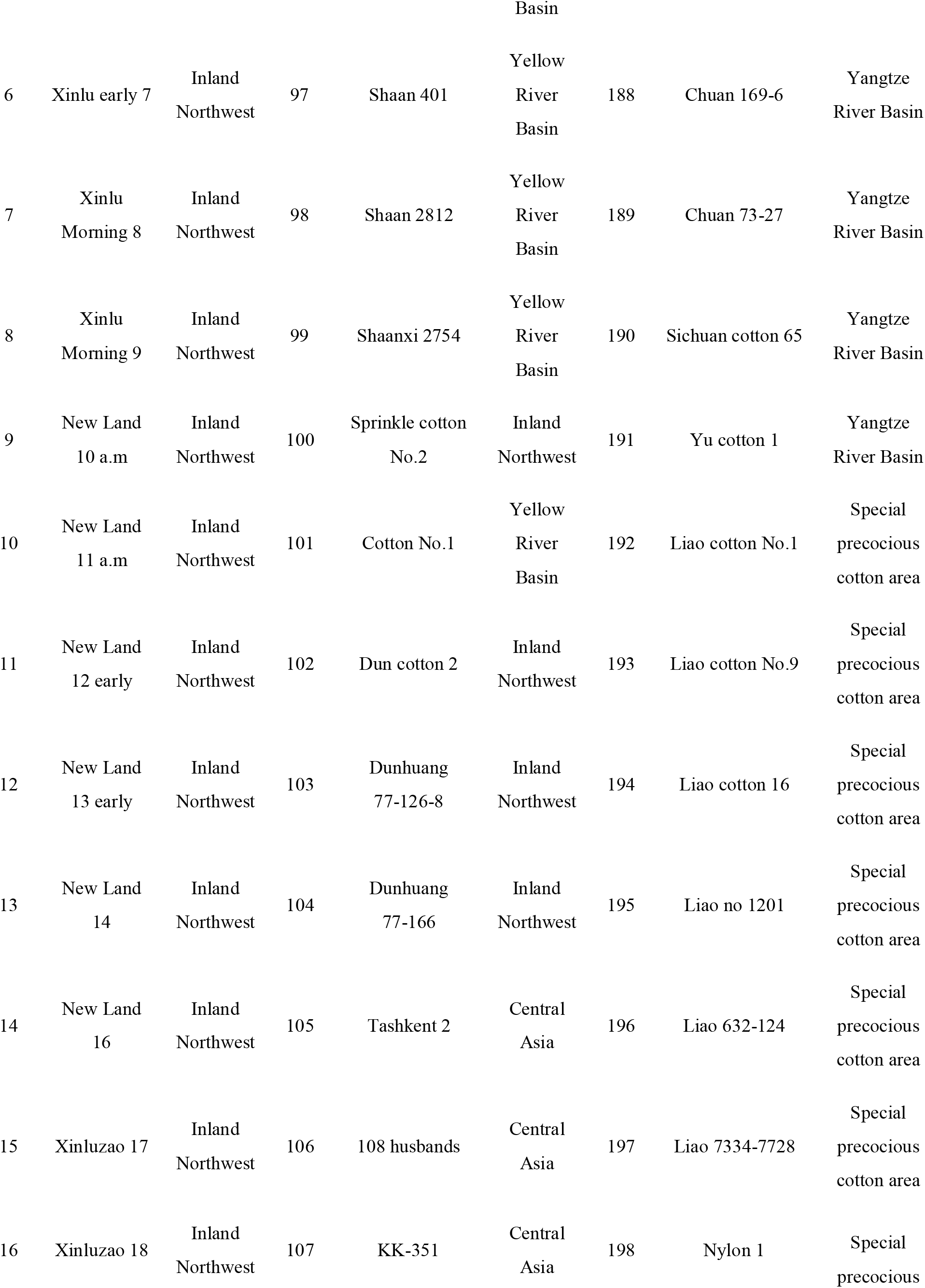
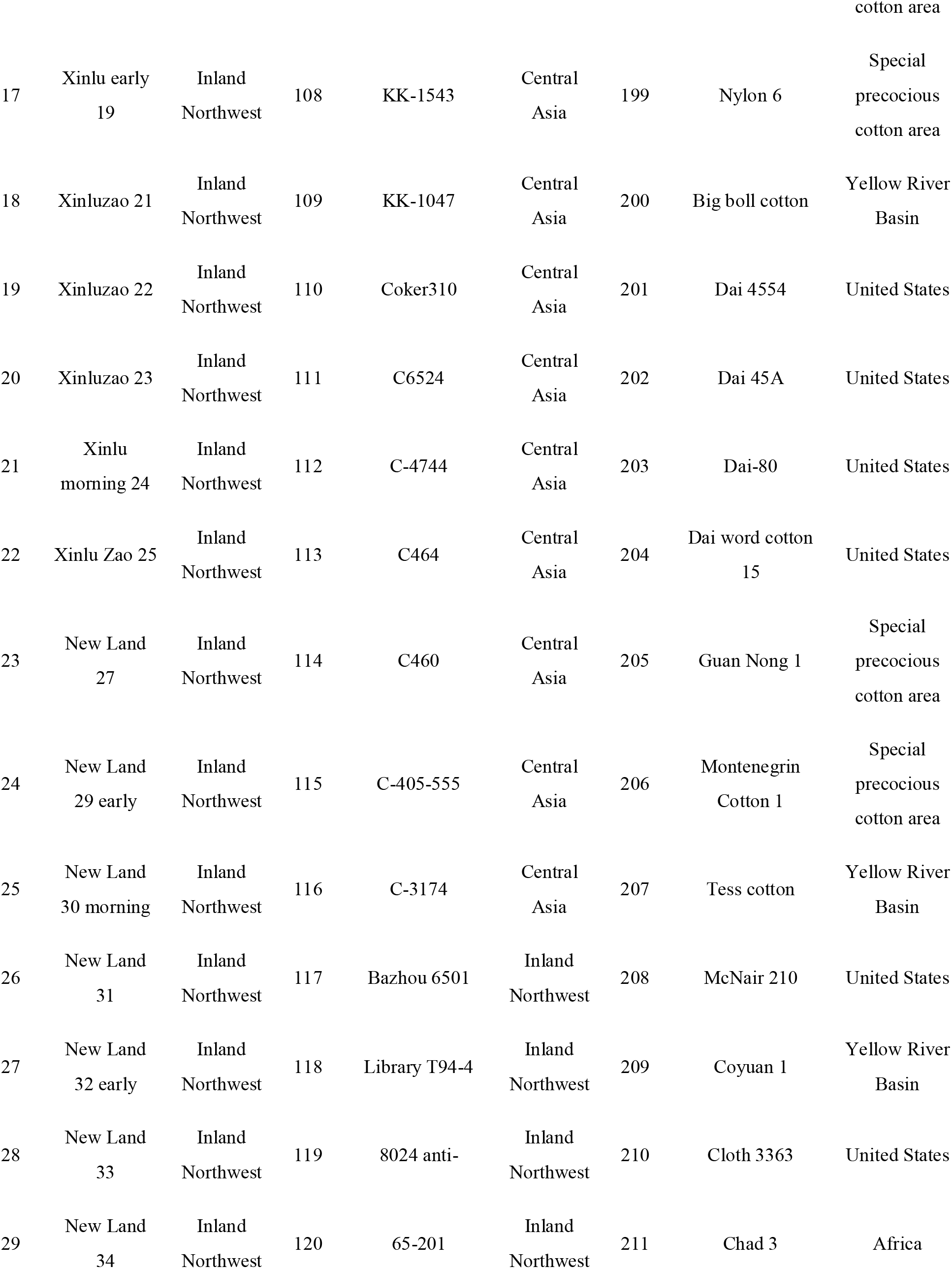
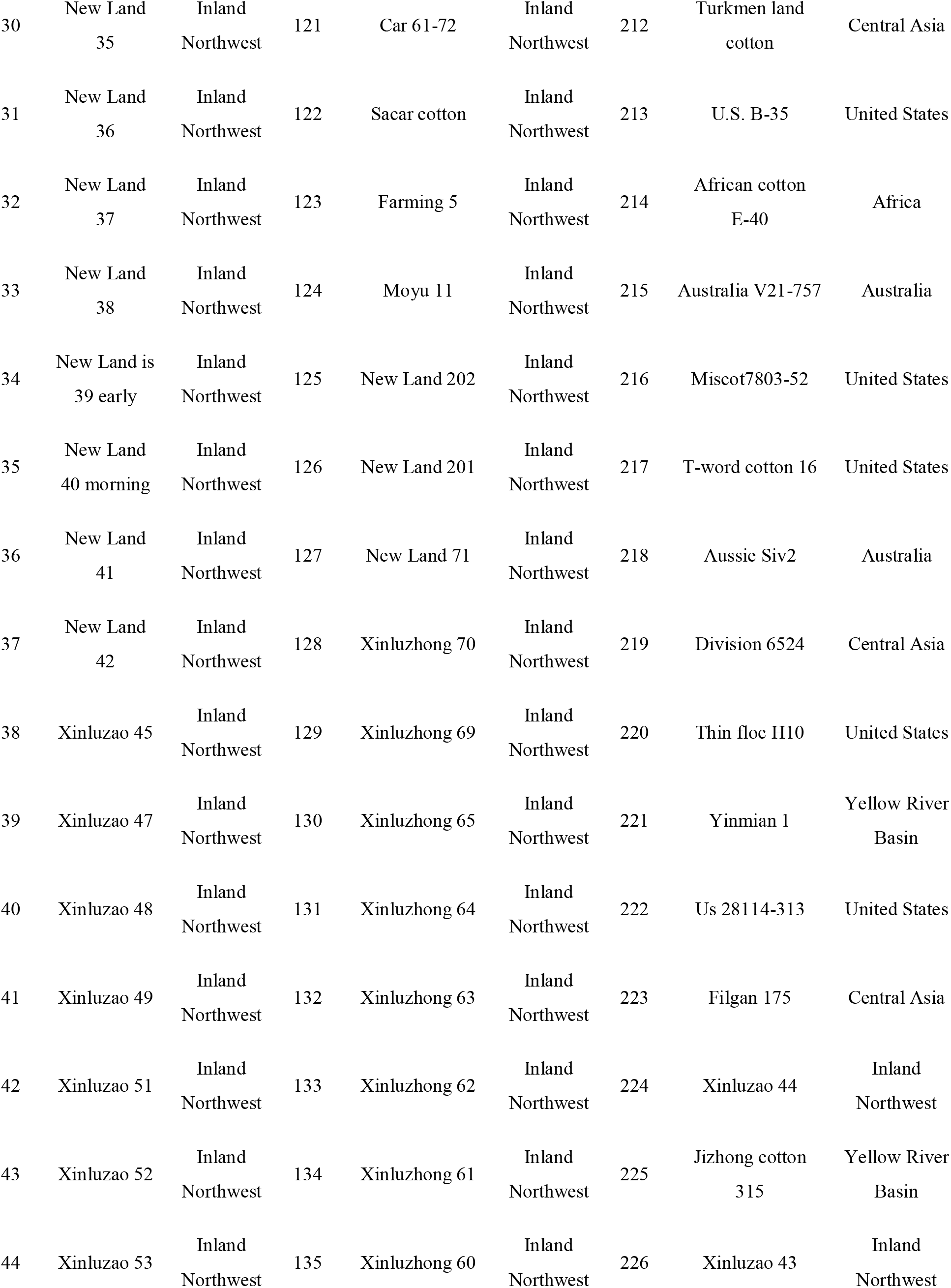
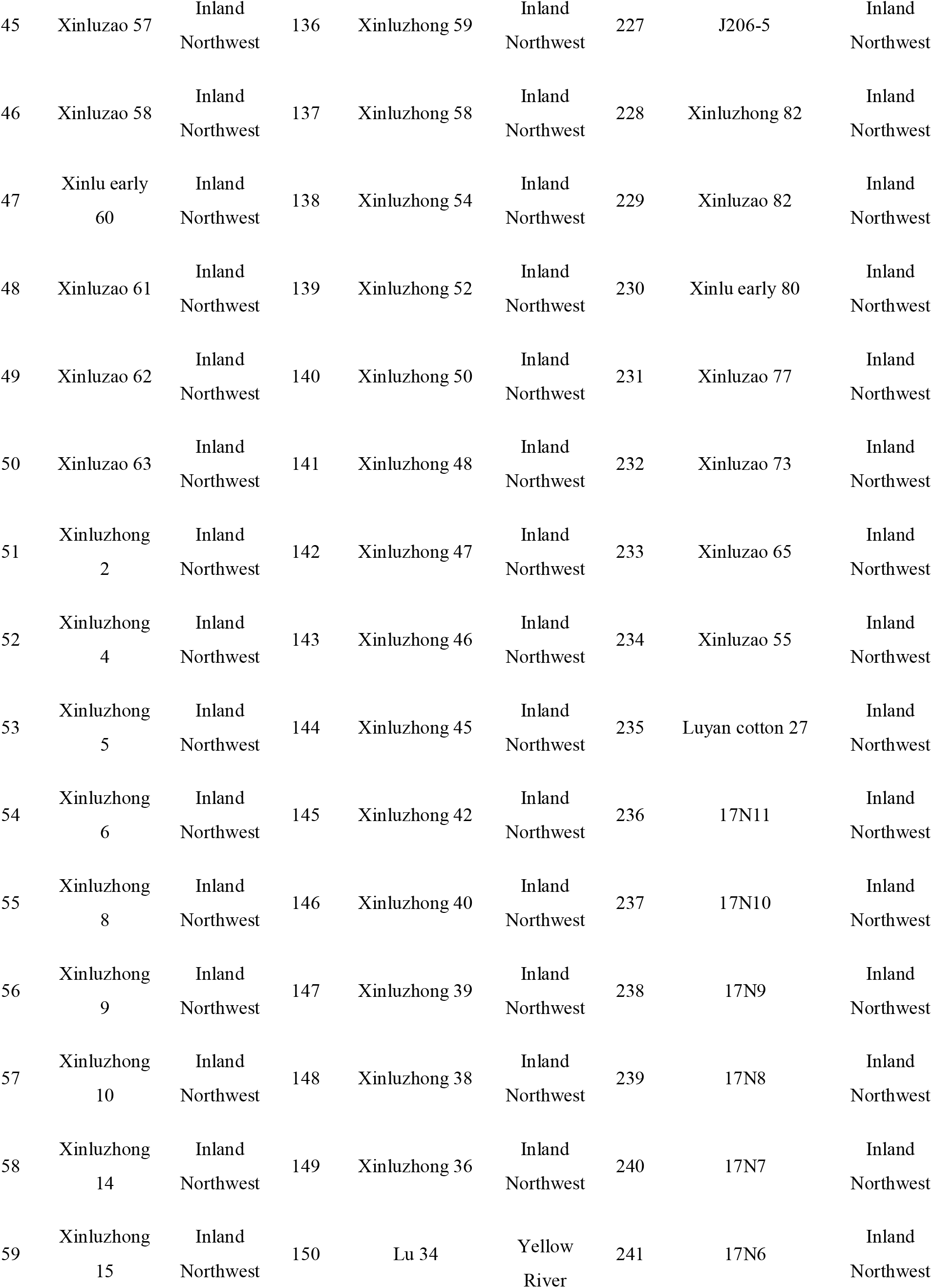
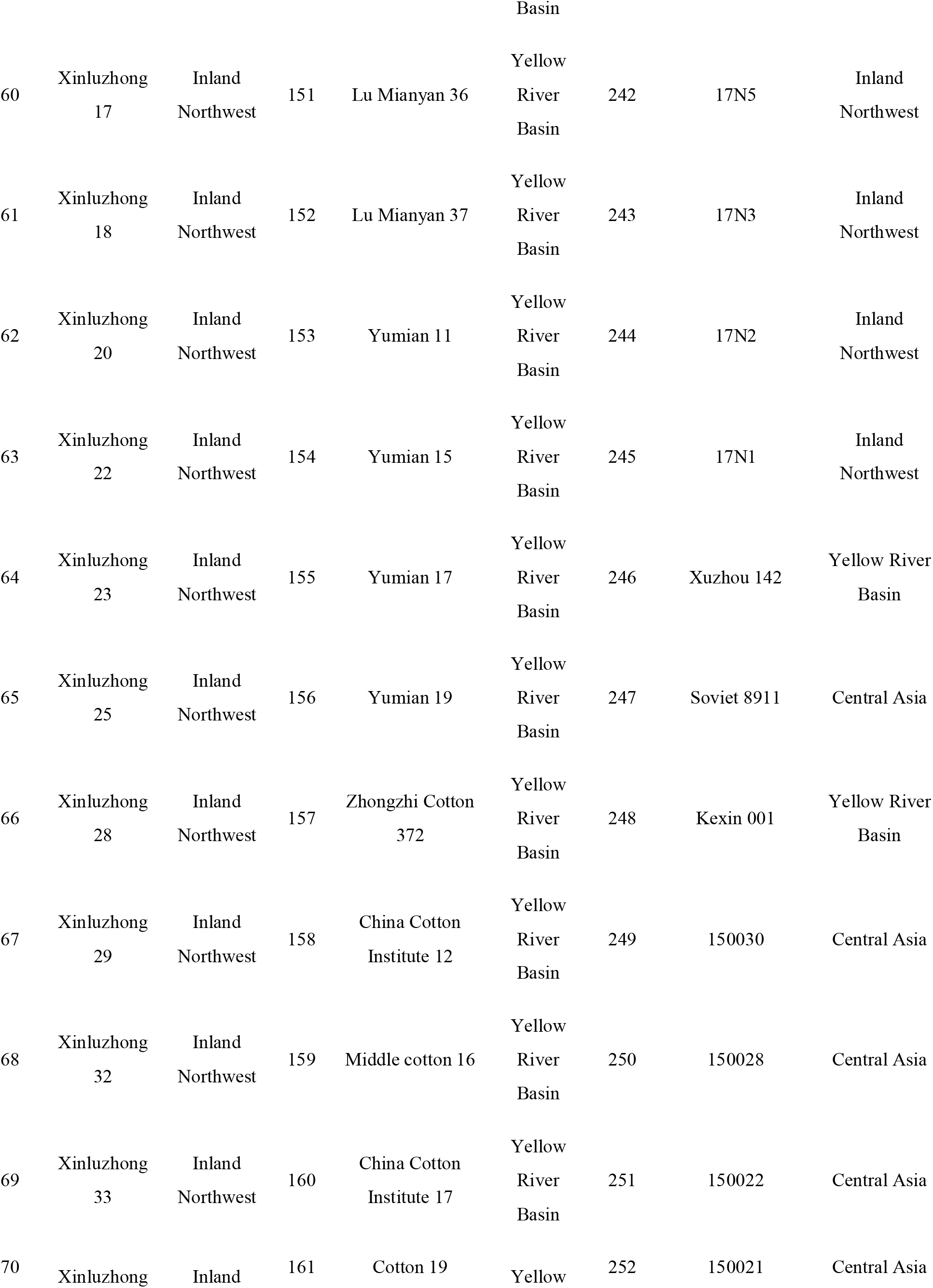
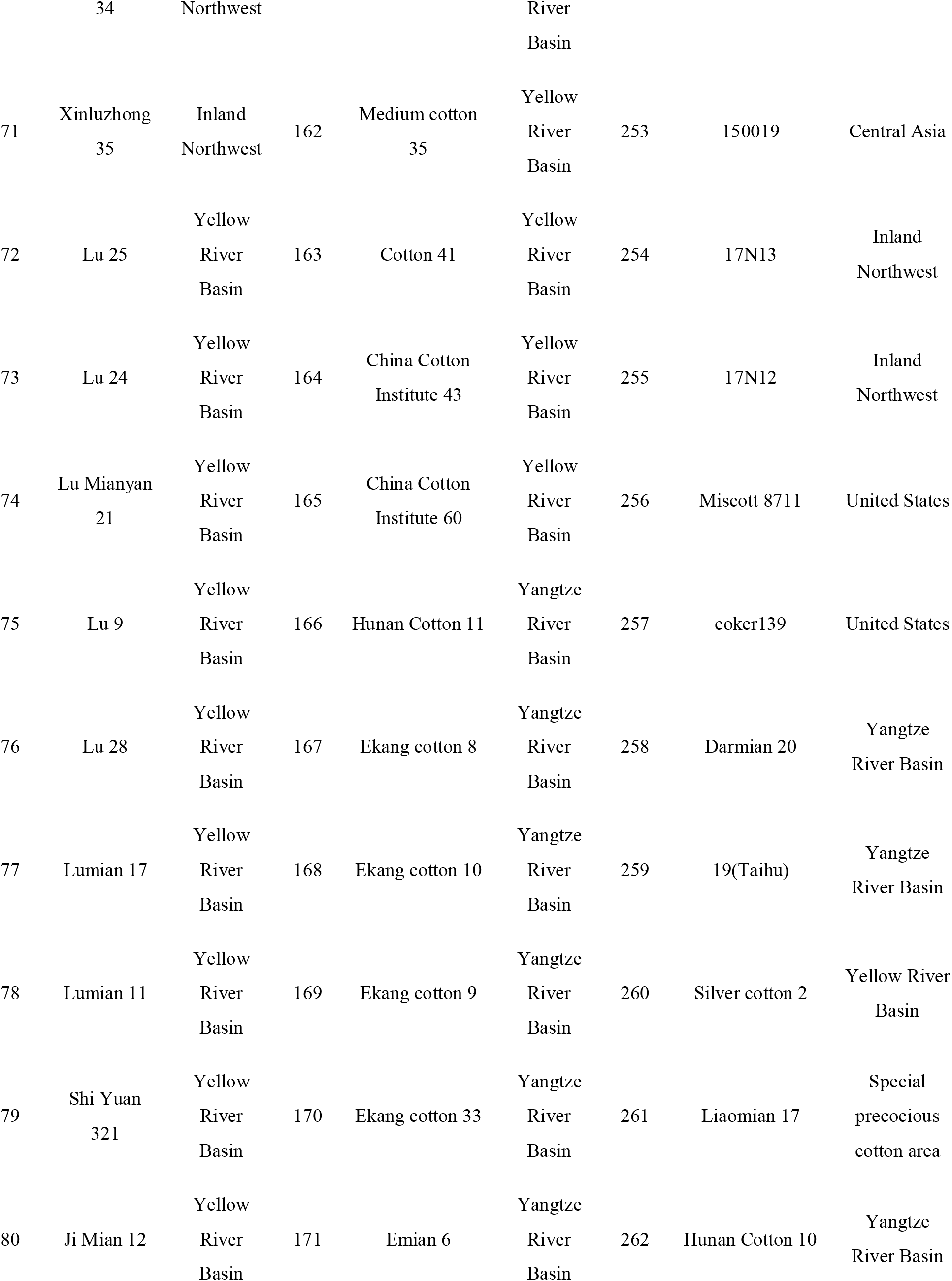
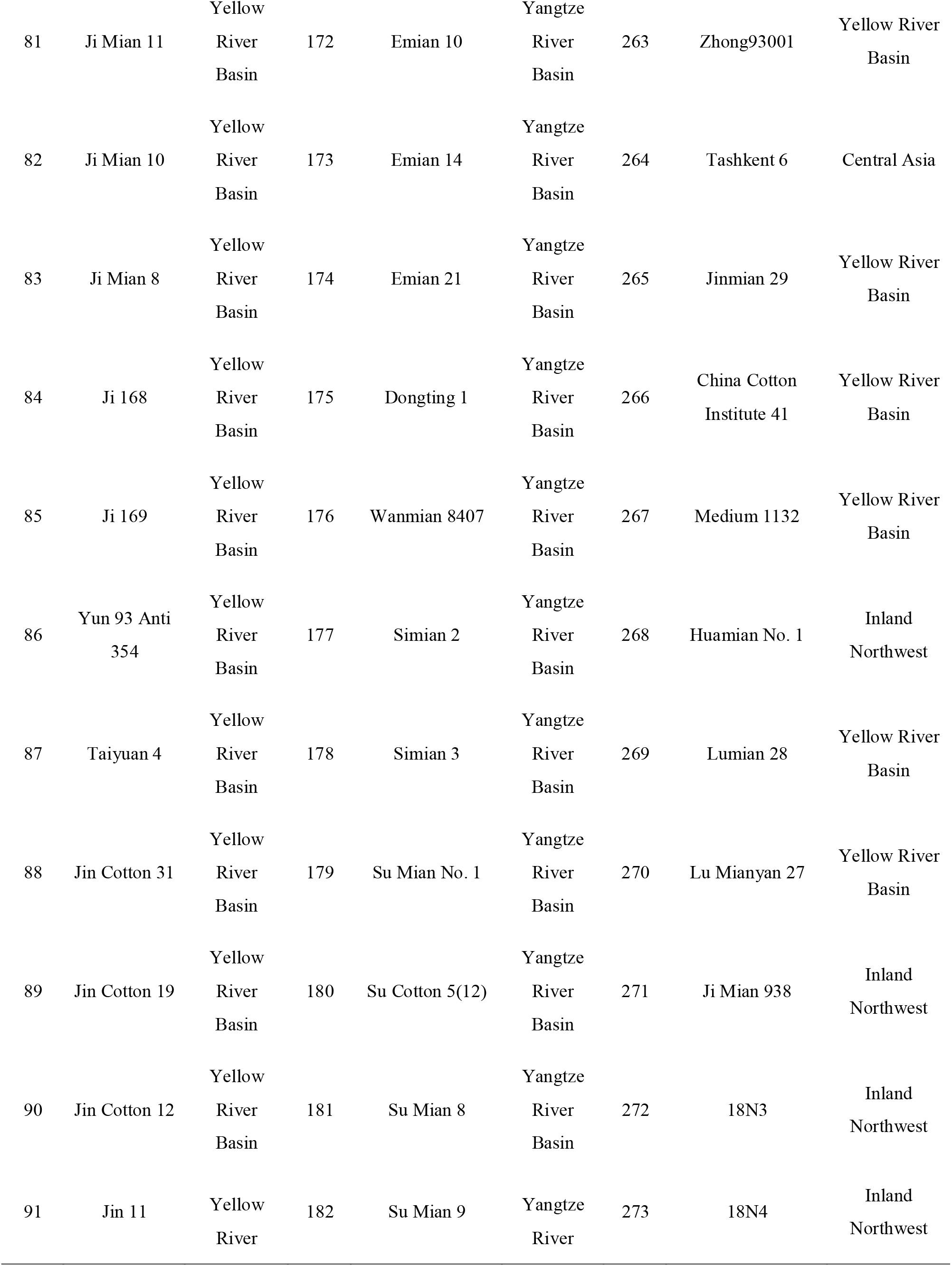
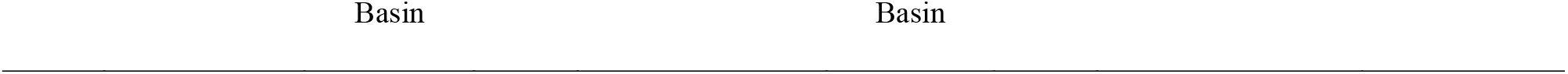
273 cotton materials.

**Appendix 2.** Statistical Table of Sample SNP Information.

| Sample ID | SNP num | Integrity | Heter ratio | Sample ID | SNP num | Integrity | Heter ratio | Sample ID | SNP num | Integrity | Heter ratio |
| --- | --- | --- | --- | --- | --- | --- | --- | --- | --- | --- | --- |
| 1 | 434,940 | 33.12% | 1.98% | 92 | 398,290 | 30.33% | 6.63% | 183 | 372,651 | 28.37% | 4.04% |
| 2 | 404,396 | 30.79% | 1.75% | 93 | 355,575 | 27.07% | 9.44% | 184 | 366,593 | 27.91% | 1.53% |
| 3 | 346,050 | 26.35% | 1.69% | 94 | 360,834 | 27.47% | 2.69% | 185 | 429,669 | 32.72% | 1.81% |
| 4 | 356,469 | 27.14% | 1.88% | 95 | 346,255 | 26.36% | 7.03% | 186 | 421,258 | 32.08% | 4.02% |
| 5 | 387,519 | 29.51% | 1.82% | 96 | 402,924 | 30.68% | 4.91% | 187 | 384,530 | 29.28% | 2.90% |
| 6 | 395,712 | 30.13% | 3.31% | 97 | 400,893 | 30.52% | 6.37% | 188 | 382,419 | 29.12% | 2% |
| 7 | 419,970 | 31.98% | 7.02% | 98 | 337,635 | 25.71% | 2.25% | 189 | 425,245 | 32.38% | 5.01% |
| 8 | 344,481 | 26.23% | 1.77% | 99 | 371,512 | 28.29% | 6.45% | 190 | 438,126 | 33.36% | 1.95% |
| 9 | 368,142 | 28.03% | 5.43% | 100 | 380,645 | 28.98% | 6.25% | 191 | 419,716 | 31.96% | 1.85% |
| 10 | 359,658 | 27.39% | 5.67% | 101 | 368,512 | 28.06% | 4.76% | 192 | 371,816 | 28.31% | 4.25% |
| 11 | 406,512 | 30.95% | 1.80% | 102 | 398,505 | 30.34% | 1.97% | 193 | 381,356 | 29.04% | 1.78% |
| 12 | 428,670 | 32.64% | 4.89% | 103 | 405,013 | 30.84% | 6.31% | 194 | 422,487 | 32.17% | 4.14% |
| 13 | 407,643 | 31.04% | 2.20% | 104 | 393,738 | 29.98% | 3.17% | 195 | 395,536 | 30.12% | 1.65% |
| 14 | 351,700 | 26.78% | 1.71% | 105 | 400,240 | 30.48% | 3.45% | 196 | 401,132 | 30.54% | 3.99% |
| 15 | 362,723 | 27.62% | 6.88% | 106 | 362,316 | 27.59% | 1.59% | 197 | 416,455 | 31.71% | 1.80% |
| 16 | 346,777 | 26.40% | 1.78% | 107 | 413,228 | 31.46% | 5% | 198 | 342,310 | 26.06% | 8.13% |
| 17 | 425,605 | 32.41% | 1.78% | 108 | 405,613 | 30.88% | 4.54% | 199 | 371,032 | 28.25% | 4.48% |
| 18 | 406,179 | 30.93% | 1.76% | 109 | 359,280 | 27.36% | 3.98% | 200 | 399,414 | 30.41% | 2.07% |
| 19 | 358,576 | 27.30% | 1.62% | 110 | 371,168 | 28.26% | 2.74% | 201 | 440,223 | 33.52% | 3.33% |
| 20 | 389,335 | 29.64% | 5.48% | 111 | 398,243 | 30.32% | 1.82% | 202 | 444,033 | 33.81% | 3.98% |
| 21 | 417,327 | 31.78% | 1.72% | 112 | 398,822 | 30.37% | 2.45% | 203 | 424,247 | 32.30% | 1.76% |
| 22 | 433,431 | 33.00% | 1.83% | 113 | 420,994 | 32.06% | 7.32% | 204 | 404,954 | 30.83% | 1.66% |
| 23 | 417,988 | 31.83% | 5.37% | 114 | 398,432 | 30.34% | 3.36% | 205 | 430,315 | 32.77% | 2.27% |
| 24 | 366,549 | 27.91% | 1.74% | 115 | 364,089 | 27.72% | 1.80% | 206 | 412,801 | 31.43% | 1.82% |
| 25 | 366,817 | 27.93% | 1.68% | 116 | 345,660 | 26.32% | 2.18% | 207 | 389,324 | 29.64% | 7.63% |
| 26 | 411,664 | 31.35% | 1.65% | 117 | 371,624 | 28.30% | 1.68% | 208 | 384,411 | 29.27% | 1.65% |
| 27 | 402,222 | 30.63% | 3.76% | 118 | 386,846 | 29.46% | 6.56% | 209 | 413,280 | 31.47% | 1.52% |
| 28 | 397,127 | 30.24% | 1.75% | 119 | 330,553 | 25.17% | 3.22% | 210 | 378,617 | 28.83% | 1.67% |
| 29 | 417,399 | 31.78% | 3.95% | 120 | 378,615 | 28.83% | 8.11% | 211 | 423,858 | 32.27% | 1.98% |
| 30 | 348,578 | 26.54% | 5.54% | 121 | 380,015 | 28.94% | 3.46% | 212 | 418,915 | 31.90% | 3.91% |
| 31 | 368,995 | 28.10% | 1.72% | 122 | 406,505 | 30.95% | 3.67% | 213 | 393,101 | 29.93% | 1.53% |
| 32 | 394,821 | 30.06% | 1.71% | 123 | 408,282 | 31.09% | 5.86% | 214 | 385,355 | 29.34% | 2.35% |
| 33 | 425,061 | 32.37% | 1.92% | 124 | 366,610 | 27.91% | 11.53% | 215 | 399,485 | 30.42% | 1.63% |
| 34 | 432,987 | 32.97% | 1.83% | 125 | 384,027 | 29.24% | 4.89% | 216 | 443,888 | 33.80% | 2.26% |
| 35 | 429,669 | 32.72% | 1.78% | 126 | 400,793 | 30.52% | 5.55% | 217 | 419,395 | 31.93% | 1.78% |
| 36 | 403,553 | 30.73% | 1.54% | 127 | 392,742 | 29.90% | 2.27% | 218 | 384,402 | 29.27% | 1.59% |
| 37 | 425,604 | 32.41% | 2.05% | 128 | 408,697 | 31.12% | 2.09% | 219 | 387,147 | 29.48% | 1.60% |
| 38 | 427,422 | 32.54% | 2.36% | 129 | 412,864 | 31.44% | 4.08% | 220 | 402,916 | 30.68% | 1.52% |
| 39 | 403,257 | 30.70% | 1.85% | 130 | 362,779 | 27.62% | 1.52% | 221 | 393,220 | 29.94% | 1.68% |
| 40 | 353,197 | 26.89% | 1.58% | 131 | 354,145 | 26.97% | 1.61% | 222 | 424,769 | 32.34% | 3.63% |
| 41 | 421,349 | 32.08% | 4.34% | 132 | 381,547 | 29.05% | 1.56% | 223 | 375,430 | 28.59% | 1.48% |
| 42 | 354,833 | 27.02% | 2.77% | 133 | 348,341 | 26.52% | 1.40% | 224 | 431,499 | 32.86% | 2.49% |
| 43 | 413,331 | 31.47% | 1.71% | 134 | 384,133 | 29.25% | 2.05% | 225 | 374,640 | 28.53% | 1.61% |
| 44 | 421,040 | 32.06% | 1.69% | 135 | 377,131 | 28.72% | 1.86% | 226 | 434,324 | 33.07% | 1.86% |
| 45 | 393,095 | 29.93% | 1.65% | 136 | 359,475 | 27.37% | 2.20% | 227 | 420,923 | 32.05% | 1.77% |
| 46 | 371,182 | 28.26% | 1.81% | 137 | 414,540 | 31.56% | 1.74% | 228 | 423,923 | 32.28% | 1.72% |
| 47 | 404,044 | 30.76% | 1.66% | 138 | 375,535 | 28.59% | 1.65% | 229 | 390,694 | 29.75% | 1.82% |
| 48 | 409,550 | 31.18% | 3.06% | 139 | 377,312 | 28.73% | 7.24% | 230 | 387,537 | 29.51% | 4.50% |
| 49 | 427,854 | 32.58% | 1.88% | 140 | 379,476 | 28.89% | 6.35% | 231 | 357,978 | 27.26% | 1.61% |
| 50 | 375,473 | 28.59% | 4.94% | 141 | 350,884 | 26.72% | 1.80% | 232 | 434,914 | 33.12% | 3.50% |
| 51 | 380,337 | 28.96% | 1.77% | 142 | 382,052 | 29.09% | 1.65% | 233 | 432,206 | 32.91% | 5.68% |
| 52 | 421,356 | 32.08% | 1.78% | 143 | 408,789 | 31.13% | 2.12% | 234 | 382,565 | 29.13% | 1.74% |
| 53 | 403,303 | 30.71% | 3.28% | 144 | 382,242 | 29.10% | 1.88% | 235 | 383,969 | 29.24% | 1.55% |
| 54 | 418,254 | 31.85% | 5.45% | 145 | 380,524 | 28.97% | 6.02% | 236 | 403,449 | 30.72% | 1.61% |
| 55 | 385,327 | 29.34% | 1.68% | 146 | 417,955 | 31.82% | 4.24% | 237 | 433,385 | 33.00% | 5.49% |
| 56 | 424,481 | 32.32% | 1.95% | 147 | 350,340 | 26.68% | 1.52% | 238 | 360,771 | 27.47% | 3.74% |
| 57 | 377,631 | 28.75% | 4.39% | 148 | 407,595 | 31.04% | 2.38% | 239 | 419,835 | 31.97% | 1.98% |
| 58 | 415,037 | 31.60% | 3.20% | 149 | 382,006 | 29.09% | 1.53% | 240 | 383,080 | 29.17% | 1.50% |
| 59 | 418,553 | 31.87% | 1.75% | 150 | 376,016 | 28.63% | 4.67% | 241 | 364,755 | 27.77% | 1.62% |
| 60 | 405,432 | 30.87% | 1.57% | 151 | 371,336 | 28.27% | 4.18% | 242 | 307,038 | 23.38% | 1.49% |
| 61 | 377,245 | 28.72% | 2.08% | 152 | 373,996 | 28.48% | 1.50% | 243 | 397,387 | 30.26% | 1.53% |
| 62 | 378,597 | 28.83% | 8.20% | 153 | 340,280 | 25.91% | 1.53% | 244 | 425,251 | 32.38% | 1.74% |
| 63 | 370,030 | 28.17% | 4.12% | 154 | 386,129 | 29.40% | 6.10% | 245 | 386,212 | 29.41% | 1.62% |
| 64 | 400,675 | 30.51% | 2.18% | 155 | 365,701 | 27.85% | 1.31% | 246 | 360,541 | 27.45% | 1.49% |
| 65 | 426,418 | 32.47% | 5.97% | 156 | 381,831 | 29.07% | 1.59% | 247 | 411,842 | 31.36% | 1.59% |
| 66 | 375,172 | 28.57% | 3.84% | 157 | 375,794 | 28.61% | 1.51% | 248 | 405,759 | 30.90% | 1.67% |
| 67 | 373,745 | 28.46% | 3.84% | 158 | 359,096 | 27.34% | 7.09% | 249 | 407,020 | 30.99% | 1.71% |
| 68 | 383,177 | 29.18% | 1.58% | 159 | 386,452 | 29.43% | 6.24% | 250 | 370,818 | 28.23% | 2.64% |
| 69 | 419,576 | 31.95% | 3.94% | 160 | 433,567 | 33.01% | 1.71% | 251 | 341,905 | 26.03% | 1.34% |
| 70 | 343,639 | 26.17% | 7.17% | 161 | 412,427 | 31.40% | 1.48% | 252 | 392,718 | 29.90% | 1.67% |
| 71 | 405,923 | 30.91% | 5.93% | 162 | 367,058 | 27.95% | 1.57% | 253 | 406,359 | 30.94% | 4.76% |
| 72 | 366,423 | 27.90% | 3.67% | 163 | 366,890 | 27.94% | 5.07% | 254 | 412,915 | 31.44% | 2.24% |
| 73 | 364,232 | 27.73% | 5.05% | 164 | 392,907 | 29.92% | 1.55% | 255 | 372,600 | 28.37% | 2.51% |
| 74 | 432,997 | 32.97% | 7.68% | 165 | 357,611 | 27.23% | 1.49% | 256 | 389,704 | 29.67% | 2.90% |
| 75 | 391,367 | 29.80% | 3.10% | 166 | 401,102 | 30.54% | 5.18% | 257 | 376,592 | 28.67% | 1.49% |
| 76 | 405,920 | 30.91% | 1.80% | 167 | 406,770 | 30.97% | 5.97% | 258 | 409,829 | 31.21% | 1.64% |
| 77 | 359,165 | 27.35% | 1.65% | 168 | 416,563 | 31.72% | 3.03% | 259 | 386,984 | 29.47% | 1.49% |
| 78 | 440,316 | 33.53% | 3.35% | 169 | 431,051 | 32.82% | 4.15% | 260 | 416,587 | 31.72% | 3.17% |
| 79 | 399,286 | 30.40% | 1.68% | 170 | 416,577 | 31.72% | 1.90% | 261 | 360,637 | 27.46% | 7.74% |
| 80 | 395,603 | 30.12% | 4.81% | 171 | 355,087 | 27.04% | 5.14% | 262 | 354,067 | 26.96% | 1.47% |
| 81 | 408,905 | 31.13% | 1.89% | 172 | 349,741 | 26.63% | 6% | 263 | 336,657 | 25.63% | 1.50% |
| 82 | 344,019 | 26.19% | 4.78% | 173 | 397,827 | 30.29% | 2.33% | 264 | 401,962 | 30.61% | 2.87% |
| 83 | 357,174 | 27.20% | 1.32% | 174 | 520,267 | 39.61% | 3.57% | 265 | 386,537 | 29.43% | 3.33% |
| 84 | 397,512 | 30.27% | 3.32% | 175 | 441,371 | 33.61% | 2.28% | 266 | 344,766 | 26.25% | 1.93% |
| 85 | 401,092 | 30.54% | 1.61% | 176 | 357,995 | 27.26% | 1.58% | 267 | 322,448 | 24.55% | 1.63% |
| 86 | 402,273 | 30.63% | 7.84% | 177 | 397,769 | 30.29% | 6.66% | 268 | 343,393 | 26.15% | 1.57% |
| 87 | 404,460 | 30.80% | 1.59% | 178 | 376,710 | 28.68% | 4.97% | 269 | 385,205 | 29.33% | 2.72% |
| 88 | 391,250 | 29.79% | 2.30% | 179 | 417,191 | 31.77% | 2.31% | 270 | 412,979 | 31.45% | 1.58% |
| 89 | 364,635 | 27.76% | 4.41% | 180 | 426,798 | 32.50% | 1.86% | 271 | 396,283 | 30.17% | 2.29% |
| 90 | 389,678 | 29.67% | 1.97% | 181 | 405,897 | 30.91% | 3.46% | 272 | 360,236 | 27.43% | 1.67% |
| 91 | 399,869 | 30.45% | 8.46% | 182 | 352,575 | 26.85% | 5.81% | 273 | 404,182 | 30.78% | 1.87% |

Note: Sample ID: sample number; SNP num: the number of SNP detected in the corresponding sample; Integrity:
the integrity of the SNP detected in the sample; Heter ratio: the heterozygosity rate of the SNP in the sample.

## Notes

https://ngdc.cncb.ac.cn/gsa

### Competing Interest Statement

The authors have declared no competing interest.

## Literature cited

Alkes L Price, Nick J Patterson, et al. 2006. Principal components analysis corrects for stratification in genome-wide association studies. Nature Genetics. 38(8), 904–909. doi:10.1038/ng1847.

Alexander D H, Novembre J, Lange K. 2009. Fast model-based estimation of ancestry in unrelated individuals. Genome research. 19(9): 1655–1664. doi:10.1101/gr.094052.109.

Ai X-T, Li X-Y, et al. 2010. Genetic Diversities of Upland Cotton Varieties in South Xinjiang. Cotton Science. 6:603–610.

Ai X-T. 2017. Genome-wide Association Analysis on Yield and Fiber Quality Traits in Upland Cotton. Xinjiang Agricultural University.

Beasley, J. O. 1940. The production of polyploids in gossypium. 31(1): 39–48.

Chen G, Du X-M. 2006. Genetic Diversity of Source Germplasm of Upland Cotton in China as Determined by SSR Marker Analysis. Journal of Genetics and Genomics. 33: 733–745.

Cingolani P, Platts A, Wang le L, et al. 2012. A program for annotating and predicting the effects of single nucleotide polymorphisms, SnpEff: SNPs in the genome of Drosophila melanogaster strain w1118; iso-2; iso-3., Fly (Austin). Apr-Jun;6(2):80–92. doi:10.4161/fly.19695.

Curt.L., Brubaker, Jonathan, F.Wendel, F Jonathan. 1994. Reevaluating the Origin of Domesticated Cotton (Gossypium hirsutum; Malvaceae) Using Nuclear Restriction Fragment Length Polymorphisms (RFLPs). American Journal of Botany. doi:10.2307/2445407.

Dong W. 2007. Genetic Diversity and SSR Abundance Analysis of Cotton Germplasm Resources. Master’s Thesis of Chinese Academy of Agricultural Sciences.

Fryxell PA. 1979. The Natural History of the Cotton Tribe (Malvaceae, Tribe gossypieae). Texas A&M University Press. p 4–7.

Fryxell PA. 1992. A Revised Taxonomic Interpretation of Gossypium L. (Malvaceae). Rheedea. 2:108–165.

Fang DD, Hinze LL, Percy RG, et al.2013.A microsatellite-based genome-wide analy sis of genetic diversity and linkage disequilibrium in Upland cotton (Gossypium hirsutum L.) cultivars from major cotton-growing countries. Euphytica. 191: 391~401. doi:10.1007/s10681-013-0886-2.

Grover CE, Gallagher JP, Jareczek JJ, Page JT, Udall JA, Gore MA, Wendel JF. 2015. Re-evaluating the Phylogeny of Allopolyploid Gossypium L. Molecular Phylogenetics and Evolution. 92:45–52. doi:10.1016/j.ympev.2015.05.023.

Gallagher JP, Grover CE, Rex K, Moran M, Wendel JF. 2017. A New Species of Cotton from Wake Atoll. Gossypium stephensii (Malvaceae). Systematic Botany. 42:115–123. doi :10.1600/036364417x694593.

Gao W, Liu F, Li S-H, Wang K-B, et al. 2010. Genetic Diversity of Allotetraploid Cotton Based on SSR Markers. ACTA AGRONOMICA SINICA. 36(11): 1902–1909.

Hardy O J, Vekemans X. 2002. SPAGeDi: a versatile computer program to analyse spatial genetic structure at the individual or population levels. Molecular ecology notes. 2(4): 618–620. doi:10.1046/j.1471-8286.2002.00305.x.

Kuang M, Yang W-H, Xu H-X. 2011. Construction of DNA Fingerprinting and Analysis of Genetic Diversity with SSR Markers for Cotton Major Cultivars in China. China Agricultural Sciences.44(1):20–27.doi:10.3864/j.issn.0578-1752.2011.01.003.

Kozich J J, Westcott S L, Baxter N T, et al. 2013. Development of a dual-index sequencing strategy and curation pipeline for analyzing amplicon sequence data on the MiSeqIllumina sequencing platform. Applied and environmental microbiology. 79(17): 5112–5120. doi:10.1128/aem.01043-13.

Liu Q. 2015. Identification of red star grass cotton-Australian cotton diploid and MSAP detection of genomic DNA methylation. Nanjing Agricultural University.

Liu W-X, Kong F-L, Guo Z-L, et al. 2003. Molecular Marker Analysis of Cotton Seed Inheritance in China since the Founding of the People’s Republic of China. Journal of Genetics and Genomics. 30(6):560–570.

Li H, Durbin R. 2009. Fast and accurate short read alignment with Burrows-Wheeler transform. Bioinformatics. 25(14):1754–1760.doi:10.1093/bioinformatics/btp324.

Li H, Handsaker B, Wysoker A, et al. 2009. The Sequence Alignment/Map Format andSAMtools.Bioinformatic.25(16):2078–2079.doi:10.1093/bioinformatics/btp352.

Multanid S,Lyon B R.1995. Genetie Finger Printing of Australian Cotton Culivars w ith RAPD Markers. Genome. 38: 1005 - 1008. doi:10.1139/g95-132.

McKenna A, Hanna M, Banks E, et al. 2010. The Genome Analysis Toolkit: a MapReduce framework for analyzing next-generation DNA sequencing data. Genome research. 20(9): 1297–1303. doi:10.1101/gr.107524.110.

Mei H-X. 2012. Genetic Diversity and Association Analysis of Main Breeding Target Traits in Uplang Cotton Cultivars of China. Nanjing Agricultural University.

Tyagi P, Gore M A, Bowman D T, et al. 2014. Genetic diversity and population struc ture in America Upland cotton (Gossypium hirsutum L.). Theoretical and Applie d Genetics. 127: 283~ 295. doi:10.1007/s00122-013-2217-3.

Wu Y-T, Zhang T-Z, Yin J-M. 2001. An Analysis about Genetic Basis of Cotton Cultivars in China since 1949 with Molecular Markers. Journal of Genetics and Genomics. 28 (11): 1040–1050.

Wu L-Y. 2012. Genetic Diversity Analysis of Sea-Island Cotton Germplasm Resources and Studies on Heterosis Utilisation between Upland Cotton and Island Cotton in the South China cotton region. Guangxi University.

Wu M, Wang N, Lin Z-X, et al. 2019. Development and evaluation of InDel markers in cotton based on whole-genome re-sequencing data. ACTA AGRONOMICA SINICA. 45(2).

Ye G-X. 2011. Genetic Analysis of cotton Cultivars Evolution in China. Nanjing Agricultural University.

